# Gli2 Overexpression Alters the Differentiation Status of Dedifferentiated Liposarcoma Cells and Results in an Immunosuppressive Myeloid Phenotype in Orthotopic Tumors

**DOI:** 10.1101/2024.09.18.613793

**Authors:** Natalie E. Bennett, Erik P. Beadle, Dominique V. Parker, Erykah J. Coe, Matthew A. Cottam, Jennifer E. Baum, Jade S. Miller, Joseph A. Serrenho, Julie A. Rhoades

## Abstract

Sarcomas are a rare classification of tumor derived from tissues of mesenchymal origin including bone, fat, muscle, cartilage, and blood vessels. These tumors often grow rapidly and have limited treatment options with few significant therapeutic advances in recent years. Liposarcomas (LPSs), the most common type of malignant soft tissue sarcoma, are derived from mesenchymal progenitors that have undergone an adipogenic lineage commitment compared to their multipotent counterparts. Interestingly, the grade of differentiation within LPS can vary highly, and the differentiation status of these tumors can drastically affect prognosis and likelihood of metastasis, making tumor differentiation a potential mechanism to target in liposarcoma development. Here, we show that overexpression of the Hedgehog transcription factor Gli2 in dedifferentiated liposarcoma (DDLPS) cells represses adipogenic differentiation while simultaneously activating markers of osteoblast differentiation *in vitro*. In addition, we observed marked differences in cytokine expression and secretion, prompting us to perform orthotopic fat pad injections of control and Gli2 overexpressing DDLPS cells. Using flow cytometry, we observed distinct changes in fat pad macrophage populations, with a particular increase in M2-like macrophages. Taken together, we find that overexpression of Gli2 in DDLPS cells alters their differentiation capacity and interactions between tumor cells and macrophages, highlighting a novel role for this developmental transcription factor in liposarcoma pathogenesis.

## Introduction

Liposarcomas are rare neoplasms of the mesenchymal stem cell (MSC) lineage that originate in adipose tissues, particularly in the retroperitoneal region, and can present in a variety of histological subtypes[1–3]. The differentiation status of liposarcomas is often considered during diagnosis, as tumors that have reverted to a more mesenchymal-like state are notably more aggressive and are associated with a worse prognosis[3]. In fact, patients with dedifferentiated liposarcoma (DDLPS) have a six-fold increased risk of death compared to patients with well-differentiated liposarcoma (WDLPS)[4].

While the exact mechanism is unknown, it is hypothesized that liposarcoma tumors that display greater levels of dedifferentiation revert to more stem cell-like phenotype allowing for greater resistance to common clinical therapies and the ability to elude immune system recognition[5]. The latter is often achieved through loss of cell surface antigens such as human leukocyte antigens (HLA), a common phenotype of sarcoma tumor-initiating cells and mesenchymal stem cells[5,6].

In our previous work, we performed bioinformatic analyses on several sarcoma datasets and found that Hedgehog signaling was enriched in DDLPS tumors, and DDLPS tumors exhibited a progenitor-like phenotype. Additionally, increased Gli2 expression correlated with gene signatures indicative of reduced immune cell infiltration in DDLPS tumors[7].

In this work, we sought to better understand the mechanism by which Gli2 may be inducing this immune-exclusive signature in patient tumors. We hypothesized that this could be a result of multiple mechanisms: 1) Gli2 may include an MSC-like phenotype in DDLPS cells with associated HLA downregulation, and/or 2) tumor cell-expressed Gli2 alters the expression of cytokines or other immune-modulatory factors.

Using SW872 human DDLPS cells, we overexpressed Gli2 and found that the tumor cells underwent a phenotypic shift and began to express markers indicative of an early osteoblast lineage commitment while losing expression of MSC markers. Additionally, we observed a marked change in expression and secretion of several myeloid recruitment cytokines. Finally, in an orthotopic model of DDLPS, we observed an increase in immunosuppressive myeloid populations as a result of tumor cell Gli2 overexpression.

## Materials and Methods

### Cell Culture

The human DDLPS cell line SW872 was a gift from the laboratory of Dr. Ben Park (Vanderbilt) and was maintained in DMEM (high glucose, pyruvate; Gibco) supplemented with 10% fetal bovine serum (FBS; Peak Serum) and 1% penicillin/streptomycin (Corning).

### Transfections

SW872 cells were stably transfected with the hGli2 FLAG3x plasmid or p3xFLAG-CMV-14 plasmid backbone as a control. hGli2 FLAG3x was a gift from Martin Fernandez-Zapico (Addgene plasmid # 84920; h-p://n2t.net/addgene:84920; RRID:Addgene_84920). The plasmid backbone sequence (p3xFLAG-CMV-14) was accessed from the Addgene vector database (Addgene ID: 1621) and custom-generated by Genscript. Cells were transfected using Lipofectamine 3000 (Invitrogen) using the manufacturer’s instructions. Transfected cells were subjected to antibiotic selection using Geneticin (G418; Gibco), and cells were passaged five times prior to use for experiments. Stable overexpression was verified via western blot before all major experiments.

### Western Blot

150,000 SW872 backbone or Gli2 overexpression (OE) cells were plated per well in a 6-well plate in complete DMEM and allowed to proliferate for 48 hours. Protein was isolated using Pierce RIPA Buffer (Thermo Scientific) supplemented with Halt Protease and Phosphatase Inhibitor Cocktail (Thermo Scientific). Protein concentration was quantified using Pierce BCA Protein Assay Kit (Thermo Scientific). Samples were loaded into a 4-20% Mini-PROTEAN TGX polyacrylamide gel (Bio-Rad) at a concentration of 20 µg per well and separated by SDS-PAGE before being transferred to a PVDF membrane using the iBlot 2 Dry Blotting System (Invitrogen). Membranes were blocked for two hours in 1x Tris-buffered saline containing 5% w/v bovine serum albumin. Membranes were then incubated with anti-Gli2 (1:500, Novus Biologicals) or anti-calnexin (Abcam, 1:1000) primary antibodies diluted in blocking buffer (BB) overnight at 4°C. The following day, membranes were incubated with anti-rabbit IgG HRP-linked secondary antibody (Cell Signaling Technology, 1:2000) diluted in BB at room temperature for 1 hour, and signal was developed using Clarity Western ECL Blotting Substrate (Bio-Rad). Membranes were imaged using the iBright CL1000 Imaging System (Invitrogen).

### RNA Extraction and Quantitative Real-Time PCR

RNA from SW872 backbone or Gli2 OE cells was collected using phenol (QIAzol; Qiagen)-chloroform (Fisher Scientific) isolation followed by isopropanol (Sigma-Aldrich) precipitation as per QIAzol manufacturer instructions. Complementary DNA (cDNA) was synthesized from 1 µg RNA using SuperScript VILO Master Mix (Invitrogen). Studies were performed using TaqMan Universal PCR Master Mix (Applied Biosystems) and TaqMan primers for the genes listed in **Table S1** (Thermo Fisher). Targets were run on custom TaqMan Array plates or as individual qRT-PCR experiments from the same samples and were normalized to the same endogenous control. Quantitative real-time PCR (qRT-PCR) was performed on the QuantStudio 7 Pro Real-Time PCR System (Applied Biosystems) using the following conditions: 2 minutes at 50°C, 10 minutes at 95°C, (15 seconds at 95°C, 1 minute at 60°C) × 40 cycles. Gene expression was calculated using the relative quantification method (ddCT) with 18S as an endogenous control. Genes that did not amplify in at least two of three biological replicates were considered to be “not detected (N.D.)” and were excluded from analysis for the group(s). Genes with only one replicate that did not amplify were retained in analysis with the non-amplified replicate removed for that group.

### Proliferation Assay

Proliferation assays were conducted using PrestoBlue Cell Viability Reagent (Invitrogen). SW872 backbone cells and SW872 Gli2 OE cells were plated at 1000 cells per well in a 96-well plate in complete media (DMEM + 10% FBS and 1% penicillin/streptomycin) and allowed to adhere overnight. The following day, cells were washed with phosphate-buffered saline (PBS; Gibco), and serum-free media was added. One day later (Day 0), media was once again replaced with 90 µL of complete DMEM. Four hours later, 10 µl of PrestoBlue reagent was added to each well (including media-only control wells), and absorbance was read at 570 and 600 nm after 30 minutes. Cells were washed with PBS and once again cultured with 90 ul of complete DMEM. On days 1, 3, 5, 7, and 9, 10 µl of PrestoBlue reagent was added to each well, absorbance was read after 30 minutes, and media was replaced with 90 µL of complete DMEM.

Reference-corrected absorbance was obtained by subtracting the 600 nm absorbance reference value from the 570 nm absorbance value. The average reference-corrected absorbance for the media-only wells was subtracted from each experimental well. Absorbance values were normalized by dividing the corrected absorbance of each biological replicate by the average Day 0 absorbance for the group (backbone versus Gli2 OE).

### Colony Forming Assay

SW872 backbone and Gli2 OE cells were plated at 100 cells per well in a 12-well plate and cultured in complete DMEM for 14 days. Cells were washed with PBS, fixed with ice-cold methanol for 20 minutes, and stained with crystal violet (Sigma) in methanol for 20 minutes followed by washing with deionized water. Individual colonies greater than 2 mm in diameter were manually counted.

### Adipocyte Differentiation and Oil Red O Staining & Quantification

SW872 backbone and Gli2 OE cells were plated at 60,000 cells per well in a 12-well plate and cultured in complete DMEM until 70-80% confluent. Media was then changed to commercial adipocyte differentiation media (Gibco) for a total of seven days with media replacement every two days. Cells were washed with PBS and fixed with 10% formalin for one hour at room temperature, washed with deionized water three times, and incubated Oil Red O Solution 0.36% in 60% isopropanol (MilliPore) for 15 minutes on a plate rocker. Cells were washed again with deionized water. Images were acquired using the Keyence BZ-X810 microscope to evaluate lipid droplet formation and cell morphology at 10X magnification. Oil Red O stain was extracted using 60% isopropanol for 15 minutes. The extracted stain was transferred to a 96-well plate, and absorbance was read at 540 nm. Protein was collected from the cells as described above, and protein concentration was quantified using the Pierce BCA Protein Assay Kit (Thermo Scientific). Oil Red O extraction value was normalized to total protein in the sample, and fold change compared to backbone control within each replicate was calculated.

### Bulk RNA Sequencing and Analysis

RNA was extracted SW872 backbone and Gli2 OE cells as described above, and samples were submitted to the Vanderbilt Technologies for Advanced Genomics (VANTAGE) core laboratory for next-generation sequencing. Briefly, mRNA enrichment and cDNA library preparation was performed using the stranded (polyA) mRNA library preparation kit (New England Biolabs), and paired-end 150 base pair sequencing was performed on the Illumina NovaSeq 6000 with an average of 50 million reads per sample. Reads from bulk RNAseq data were processed using the nf-core/RNAseq pipeline[8]. Aligned transcript counts were imported into R v4.2.2 using Tximport v1.32.0[9]. Low capture genes with less than ten total counts summed across all samples were removed. Differential expression was performed with DESeq2[10]. Genes were considered to be significantly different if the log2 fold change was greater than 1 and adjusted p value was less than 0.05. Functional enrichment analysis was performed with MetaScape v3.5.20240101[11] to evaluate enrichment of Hallmark, GObp, KEGG, and Reactome gene sets.

### Cytokine Quantification

150,000 SW872 backbone and Gli2 OE cells per well were plated in a 6-well dish and cultured for 48 hours. Conditioned media was collected and centrifuged at 1000 x g for 10 minutes. Supernatant was collected and stored at -80°C until ready to use for cytokine assays. Samples were submitted to the VUMC Analytical Services Core for measurement of cytokines from the MILLIPLEX MAP Human Cytokine/Chemokine Magnetic Bead Panel (Millipore Sigma). The same conditioned media was used for the M-CSF (CSF-1) Human ELISA Kit (Invitrogen), which was performed using manufacturer instructions. Total protein concentration was quantified using the Pierce BCA Protein Assay (Thermo Fisher) and used to normalize cytokine values.

### Orthotopic Animal Model of Dedifferentiated Liposarcoma

Male and female athymic Nude-Foxn1^nu^ (7 weeks old, Jackson Laboratory) were anesthetized under continuous isoflurane inhalation and injected with 2x10^6^ SW872 backbone or Gli2 OE cells in sterile saline bilaterally into the dorsal/posterior aspect of the inguinal fat pad via subcutaneous route. Mice were monitored daily and weighed two times per week. Mice were euthanized 18 days after tumor injection, and both fat pads were collected and weighed.

### Flow Cytometry

100 mg of collagenase IV was resuspended in 1 ml of PBS. Diluted collagenase was added to serum-free DMEM in a 1:100 ratio. Fat pads were finely minced and placed in a 1.5 ml Eppendorf tube with 1 ml of the collagenase solution and were digested for three hours at 37°C in a tube rotator. Digested fat pads were then passed through a 70 µm filter and rinsed with complete DMEM. Cells were collected via centrifugation, and red blood cells were lysed using ACK Lysing Buffer (Gibco). 5x10^5^ cells per test were stained for viability with Ghost Dye Violet 510 (Tonbo) at a dilution of 1:5000 followed by Fc receptor blocking using a combination of 0.5 µL per test of Purified Anti-Mouse CD16/CD32 Fc block (Tonbo) and 1 µL per test of Human Trustain FcX (Biolegend). Samples and fluorescence minus one controls were stained using antibodies at the dilutions listed in **Table S2**. Single stain controls were prepared using compensation beads (Invitrogen). Samples were fixed with 2% paraformaldehyde, and flow cytometry was performed on the Cytek Aurora spectral flow cytometer (Cytek Biosciences). Analyses were completed using FlowJo software (BD Life Sciences).

### Statistics

All statistical analyses were performed using Prism 10.2.0 (GraphPad Software). Values are presented as mean and standard deviation (SD). Unpaired t-test (*p < 0.05, **p < 0.01) or multiple unpaired t-tests with correction for multiple comparisons (FDR, Benjamini-Hochberg procedure, Q = 5%, *q < 0.05) were performed for one or more than one gene or protein target, respectively. Outlier testing was performed using ROUT method (Q < 1%). *In vitro* experiments were performed with three independent biological replicates unless otherwise specified.

### Ethics Statement

All animal protocols were approved by Vanderbilt University Institutional Animal Care and Use Committee and were conducted according to National Institutes of Health guidelines for care and use of laboratory animals.

## Results

### Characterization of a Gli2-overexpressing dedifferentiated liposarcoma cell line

The SW872 liposarcoma cell line, a model of human dedifferentiated liposarcoma[12,13], was transfected with a hGli2 FLAG3x plasmid or p3xFLAG-CMV-14 plasmid backbone as a control. Stable Gli2 overexpression was confirmed via western blot after antibiotic selection and passaging (**Figure 1A**). Expression of key Hedgehog signaling genes was evaluated to confirm functional Gli2 overexpression. As expected, Gli2 overexpressing cells (Gli2 OE) had approximately thirteen-fold higher *Gli2* gene expression than backbone control (**Figure 1B**). *Gli1* and *HHIP* (Hedgehog interacting protein) were also increased in Gli2 OE cells, the former being another transcription factor in the Hedgehog pathway that is a target of Gli2[14]. Hedgehog interacting protein is an inhibitor of Hedgehog signaling but also a target of Hedgehog/Gli signaling[15]. Altogether, these results demonstrate that Gli2 overexpression does indeed increase the expression of Hedgehog pathway components.

**Figure 1.**
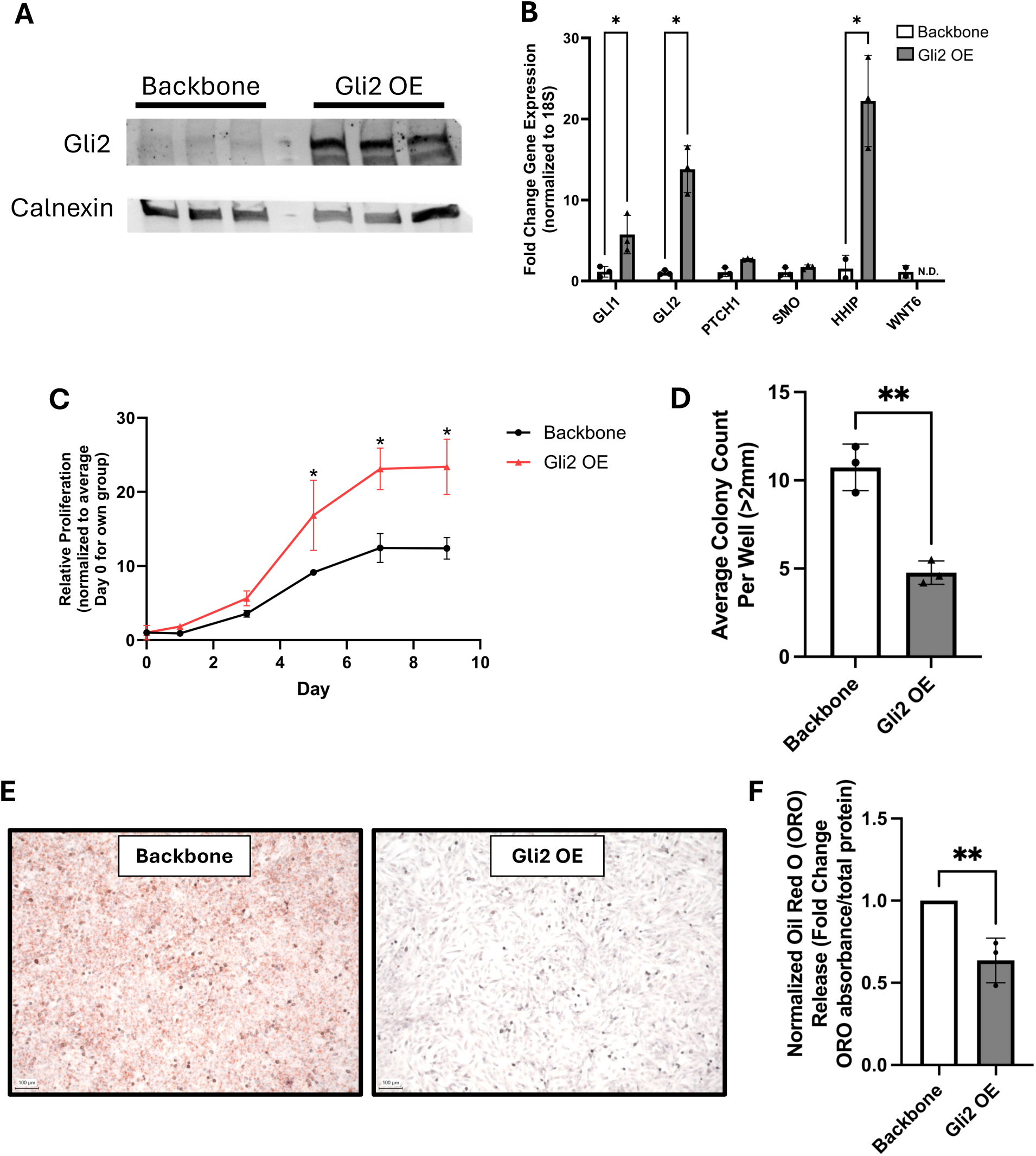
Characterization of SW872 Gli2 overexpressing cells. (A) Western blot of Gli2 and calnexin (loading control) in SW872 backbone and Gli2 OE cells. (B) Gene expression of Hedgehog signaling components by qRT-PCR. (C) Proliferation of backbone and Gli2 OE cells over time as measured by PrestoBlue assay. (D) Colony forming assay quantification of average colonies per well. (E) Representative images of SW872 backbone and Gli2 OE cells after culturing in adipocyte differentiation medium. (F) Quantification of Oil Red O release assay from cells subjected to adipocyte differentiation. Mean and standard deviation shown. *q < 0.05 **p < 0.01. Graphs represent biological replicates from three independent experiments. N.D. = not detected.

The proliferation of Gli2 OE SW872 cells was measured over the course of nine days using the PrestoBlue proliferation assay. From five days onward, Gli2 OE cells had higher proliferation compared to baseline than backbone control cells (**Figure 1C**). We hypothesized that Gli2 overexpression would induce a more mesenchymal stem cell-like phenotype in the liposarcoma cells, given the role of Hedgehog signaling in reducing adipogenesis in MSCs in non-malignant contexts[16–18]. To assess the self-renewing capacity of Gli2 OE and backbone cells, a colony forming assay was also performed. Interestingly, Gli2 OE cells had lower colony counts (**Figure 1D**; example plate in **Figure S1**), suggesting reduced ability to self-renew.

Finally, Gli2 OE and backbone cells were cultured with adipogenic differentiation media and stained with Oil Red O to evaluate the presence of lipid droplets, demonstrating a notable reduction in droplet staining in the Gli2 OE cells (**Figure 1E**). This stain was then extracted for quantification and normalized to total protein. Gli2 OE cells had an approximately 36% reduction in normalized Oil Red O staining compared to paired SW872 backbone control samples upon culturing with adipogenic differentiation media (**Figure 1F**). Therefore, Gli2 overexpression reduces the ability of these DDLPS cells to differentiate toward the adipocyte lineage. This finding, coupled with the increased proliferative capacity but reduced self-renewal capacity of Gli2 OE cells, warrants further exploration of the transcriptional pathways enriched in these cells that may contribute to their phenotype.

### Gli2-overexpressing dedifferentiated liposarcoma cells have a distinct transcriptional profile with notable differences in immunoregulatory and skeletal developmental pathways

Because of the evident differences in our initial phenotypic characterization of SW872 cells with overexpressed Gli2 compared to control, we sought to better characterize the transcriptional profile of these cells through bulk RNA sequencing. Upon quality control and read alignment, differential expression was performed using DESeq2[10], and functional analysis of significantly different genes was performed with MetaScape[11] to evaluate pathway enrichment through a variety of methods, such as Hallmark, Gobp, KEGG, and Reactome gene sets. The two cell lines were indeed transcriptionally distinct, with 937 unique downregulated genes and 1304 unique upregulated genes in the SW872 Gli2 OE cells compared to backbone control (**Figure 2A**). As expected, *GLI2* was among the genes with the highest fold increase in Gli2 OE cells. Examination of the top 20 enriched clusters from Gene Ontology and pathway enrichment analysis then showed enrichment in a number of developmental pathways, such as “GO:0001944: vasculature development,” “GO:0007423: sensory organ development,” “GO:0007507: heart development,” and “GO:0048729: tissue morphogenesis” (**Figure 2B**), all of which are reasonably expected given the role of Gli2 in embryogenesis and tissue patterning[19]. Additionally, there was notable enrichment in “M5930: HALLMARK EPITHELIAL MESENCHYMAL TRANSITION” among upregulated genes in the Gli2 OE group, underscoring the role of Gli2 in essential processes that dictate tumor progression.

**Figure 2.**
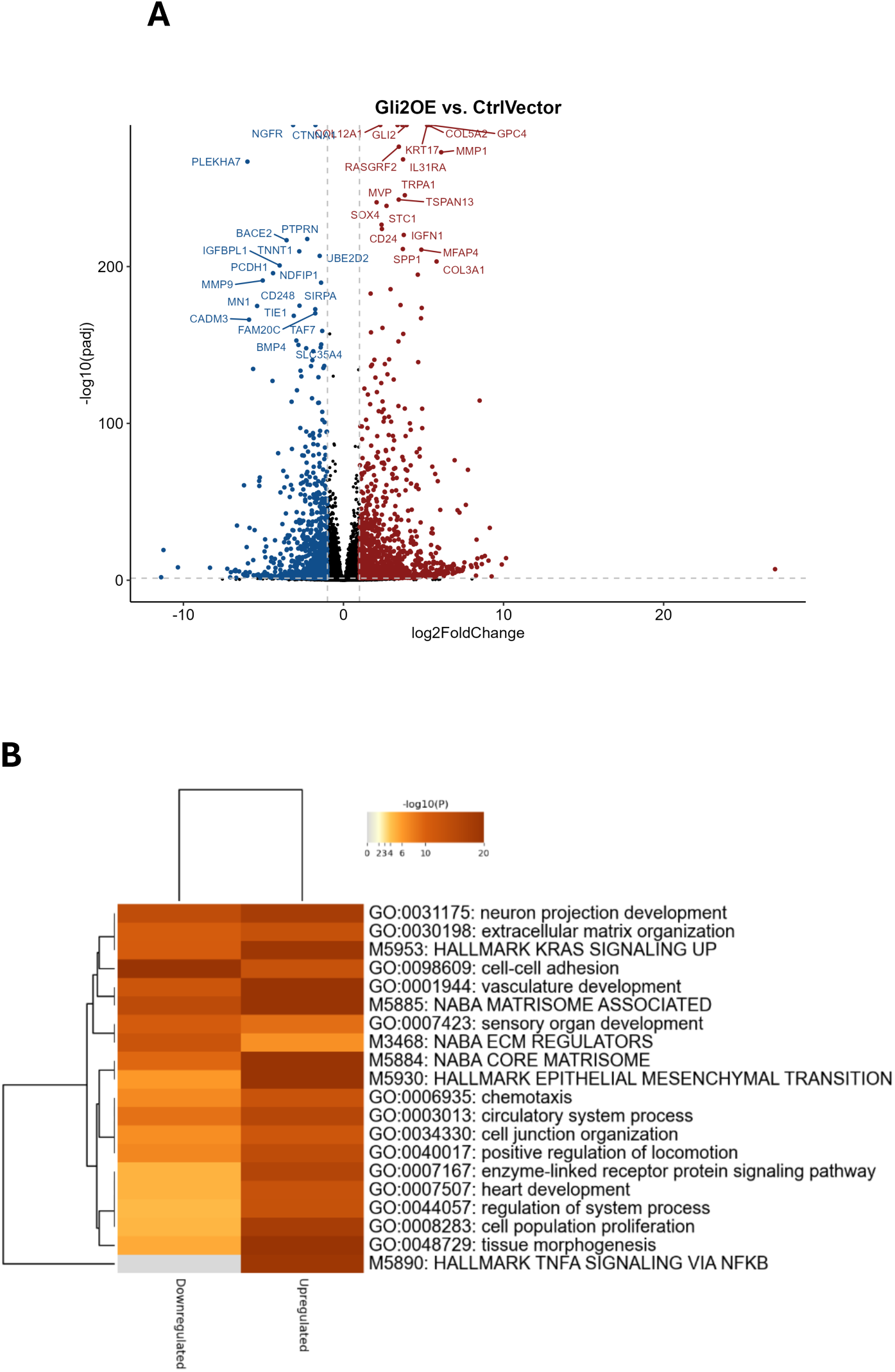
Bulk RNA sequencing reveals a distinct transcriptional profile in Gli2-overexpressing DDLPS cells. SW872 cells were transfected with either Gli2 overexpression vector or empty vector control. (A) Bulk RNA sequencing was performed, and differential genes were plotted. 937 unique downregulated genes and 1304 unique upregulated genes were identified. (B) Metascape was used to perform Gene Ontology and pathway enrichment analyses. The top 20 enriched clusters for downregulated and upregulated genes are shown. A more comprehensive heatmap can be found in supplemental information.

Expansion of analysis to the top 100 enriched clusters revealed pathways that may contribute to our prior observations of association between Gli2 and gene signatures suggestive of differential immune infiltration in publicly available sequencing databases of patient DDLPS samples[7]. Our tumor cell sequencing demonstrated differences in pathways such as “GO:0006954: inflammatory response,” “GO:0050727: regulation of inflammatory response,” “M5932: HALLMARK INFLAMMATORY RESPONSE,” “M5947: HALLMARK IL2 STAT5 SIGNALING,” and “M5890: HALLMARK TNFA SIGNALING VIA NFKB” in Gli2 OE cells versus backbone control (**Figure S2**). While these are broad characterizations, they support the notion of exploring more granular changes in specific genes that may recruit or exclude immune cells.

Of particular note was the enrichment of GO:0001501: skeletal system development” and “GO:0001503: ossification” among the upregulated genes in Gli2 OE cells (**Figure S2**), given the role of Hedgehog signaling in modulating the balance between adipocyte and osteoblast differentiation[16–18]. In fact, *SPP1*, a marker of osteoblast differentiation, was among the top upregulated genes in Gli2 OE cells (**Figure 2A**). These findings, alongside the reductions in self-renewal capacity (**Figure 1D**) and adipogenic potential (**Figure 1E-F**) in Gli2 OE cells underscore that Gli2 may play a role in dictating the differentiation fate of DDLPS tumor cells.

### Gli2-overexpressing dedifferentiated liposarcoma cells have increased expression of osteoblast markers

Prior literature suggests that Gli2 impairs MSC differentiation to the adipocyte lineage[16–18], often enhancing MSC differentiation into osteoblasts. Recent work has also demonstrated the pluripotency of patient-derived DDLPS cells when subjected to adipogenic or osteogenic media[20]. However, the role of Gli2 in maintaining the MSC-like phenotype of DDLPS cells has not been explored. Therefore, we predicted SW872 Gli2 OE cells would have decreased expression of adipocyte markers and increased expression of MSC markers compared to SW872 backbone control cells. We performed qRT-PCR for a panel of adipocyte differentiation markers (**Figure 3A**) to identify the differentiation status of Gli2 OE cells compared to backbone control. There was no significant difference in adipocyte markers, although *LPL* (lipoprotein lipase) trended downward in the Gli2 OE cells (**Figure 3B**). Neither cell line expressed *ADIPOQ* (adiponectin) or *LEP* (leptin). We next evaluated genes that are typically increased in MSCs. To our surprise, Gli2 OE cells had significantly lower expression of *THY1* and a trending decrease in *PDGFB* and *NES* (**Figure 3C**). This suggests that Gli2 overexpression may not be pushing SW872 dedifferentiated liposarcoma cells toward an MSC-like phenotype.

**Figure 3.**
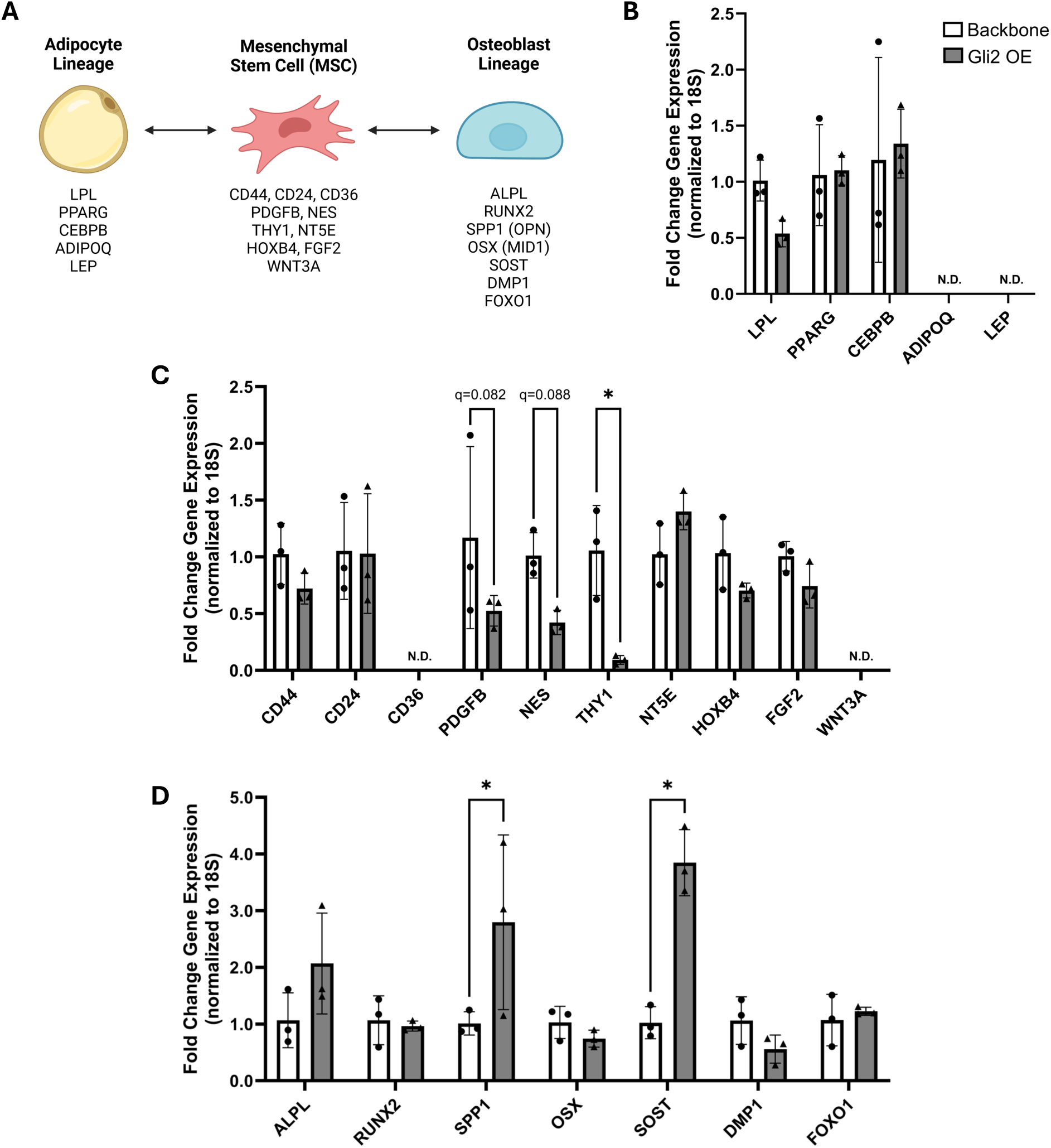
Gli2 overexpression results in differential expression of key genes representing MSC differentiation states. (A) Schematic of differentiation of MSCs to the adipocyte lineage and osteoblast lineage. Genes listed are characteristic of the cell types. Graphs show expression of (B) adipocyte lineage, (C) mesenchymal stem cell, and (D) osteoblast lineage genes in SW872 backbone and Gli2 OE cells. Mean and standard deviation shown. *q < 0.05. Biological replicates from three independent experiments. N.D. = not detected. Schematic created with BioRender.com

However, as described above, Gli2 is known to drive osteoblast differentiation from MSCs[16,21–23]. In Gli2 OE SW872 cells, there was increased expression of *SPP1* (osteopontin), a key marker of osteoblast differentiation that was noted on bulk RNA sequencing above (**Figure 2A**), and *SOST* (sclerostin), which is commonly expressed by osteocytes, a cell type terminally differentiated from osteoblasts (**Figure 3D**). *DMP1* and *FOXO1*, additional markers of osteocyte differentiation, were not increased, thereby suggesting that increased sclerostin expression in Gli2 OE cells is indicative of differentiation toward the osteoblast lineage but not terminal differentiation to osteocytes. Sclerostin has been shown to increase adipocyte differentiation[24]. However, sclerostin is expressed in skeletal sarcomas, such as osteosarcoma, and is positively correlated with osteogenic differentiation[25]. Given the known role of Gli2 in inducing osteoblast differentiation, these results suggest that rather than driving dedifferentiated liposarcoma cells toward an MSC-like phenotype, Gli2 overexpression may induce an osteoblast-like phenotype, which is further supported by the reduced self-renewal capacity shown in **Figure 1D**.

Finally, we evaluated markers of epithelial to mesenchymal transition and Notch signaling to further characterize these potential changes in differentiation state. As expected, neither cell line expressed *CDH1* (E-cadherin), but Gli2 OE cells had a significant reduction in *CDH2* (N-cadherin), further supporting that Gli2 overexpression pushes SW872 cells away from a mesenchymal-like phenotype (**Figure S3A**). N-cadherin overexpression has been shown to inhibit osteogenesis[26], so reduction of this marker in Gli2 OE cells lends additional support to the findings of increased osteoblastic lineage markers observed in **Figure 3**. Furthermore, TGF-β1 increased with Gli2 overexpression, whereas TGF-β2 decreased. However, the effect of the TGF-β superfamily on osteoblastogenesis is variable depending on phase of differentiation and activity of other signaling pathways[27–29]. Additionally, Gli2 OE cells also had increased expression of *NOTCH3*, *HEY1*, and *JAG1* (**Figure S3B**). Notch inhibition promotes adipogenesis[30], whereas Jag1/Notch signaling increases osteogenesis of human mesenchymal stem cells and inhibits adipogenesis[31]. These results demonstrate that rather than inducing a more MSC-like state in dedifferentiated liposarcoma cells, Gli2 overexpression induces a transcriptional profile more characteristic of osteoblastic differentiation.

### Gli2 overexpression results in differential expression of cytokine genes and cytokine secretion

Our previous analyses of sequencing data from patient differentiated liposarcoma tumors suggest that Gli2 expression is inversely correlated with markers of antigen presentation and myeloid and T-cell populations, suggesting that Gli2-high tumors may have reduced immune infiltrate[7]. These findings are further supported by bulk RNA sequencing of SW872 backbone and Gli2 OE cells, which showed enrichment of various immunoregulatory pathways (**Figure S2**). We therefore sought to better characterize specific genes that may contribute to these observations. As predicted based on our previous work, there was a trend toward decrease in HLA-B expression in Gli2 OE cells, and while HLA-C was expressed in backbone control cells, this MHC class I molecule was not expressed in Gli2 OE cells, suggesting a decrease in expression as well (**Figure 4A**). Additionally, CD274 (PD-L1) significantly increased in Gli2 OE cells.

**Figure 4.**
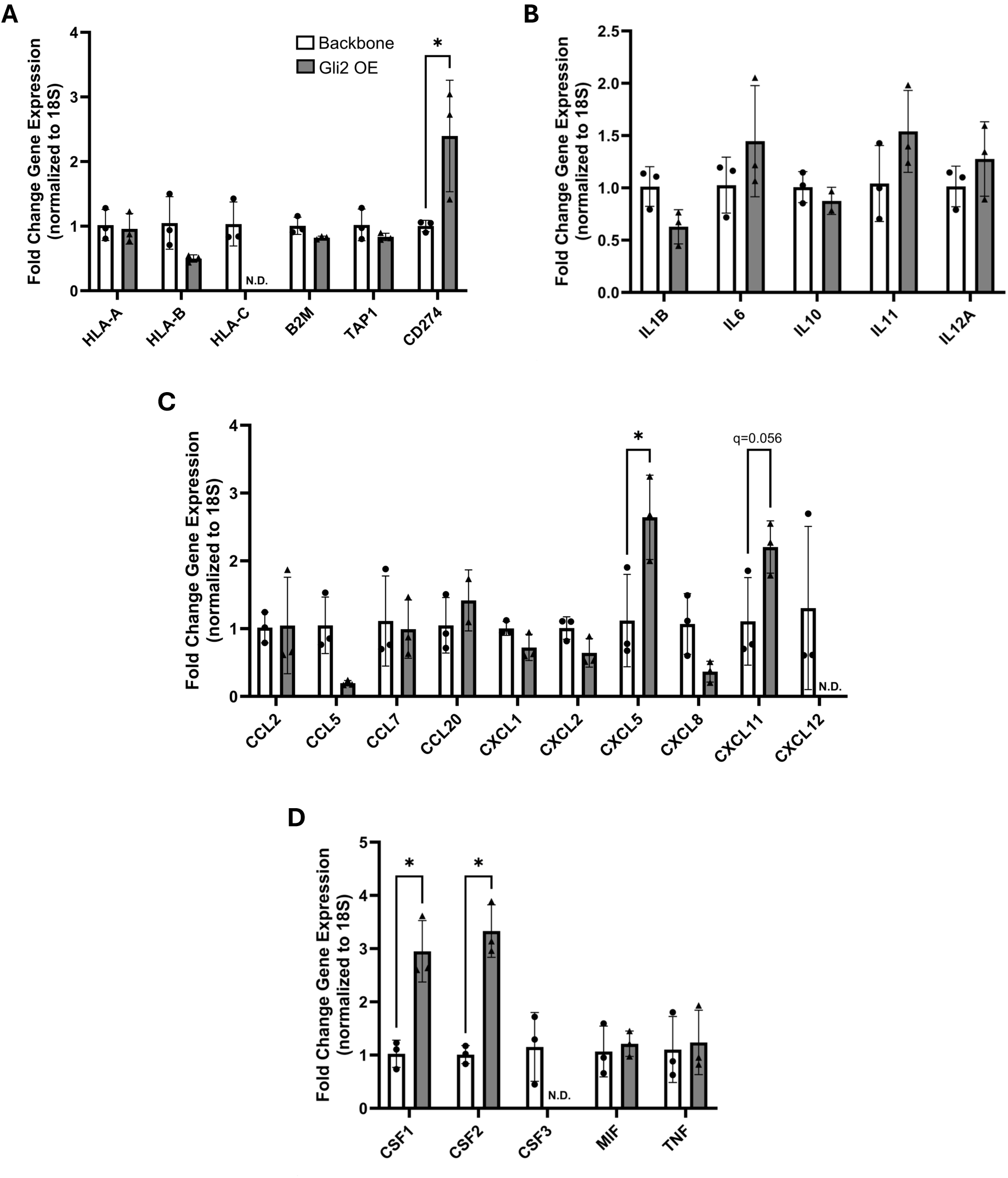
Gli2-overexpressing SW872 cells have altered expression of key immune signaling genes. Gene expression of (A) antigen presentation genes and CD274 (PD-L1), (B) interleukins, (C) chemokines, and (D) other cytokines measured by qRT-PCR. Mean and standard deviation shown. *q < 0.05. Biological replicates from three independent experiments. N.D. = not detected.

Among the list of interleukin, chemokine, and other cytokine genes assayed, there was a significant increase in *CXCL5* expression as well as an approximately three-fold increase in *CSF1* and *CSF2* expression in Gli2 OE cells (**Figures 4B-D**). Among cytokines evaluated, several did not amplify in both groups: *CCL4*, *CCL11*, *CXCL9*, *CXCL10*, *IL1A*, *IL2*, *IL4*, *IL12B*, *IL13*, *IL17A*, *IL18*, *IL23A*, and *IFNG*. Conditioned media from these cells also showed a differential secretory profile with Gli2 overexpression. In Gli2 OE cells, there was a significant decrease in CCL5 and CXCL8 (IL-8) (**Figure 5A**), consistent with trends demonstrated at the transcriptional level. Both cell lines secreted an abundance of CCL2, which is known to be secreted by adipose tissue[32]. Furthermore, an enzyme-linked immunosorbent assay (ELISA) for M-CSF (CSF-1) demonstrated higher secretion in Gli2 OE cells (**Figure 5B**), again consistent with gene expression data in **Figure 4**. In total, this data demonstrates that while reduced HLA expression at the gene level in Gli2 OE cells may be a potential mechanism for a reduced immune infiltration signature in dedifferentiated liposarcomas, there may be a role for Gli2-associated changes in cytokine expression and secretion.

**Figure 5.**
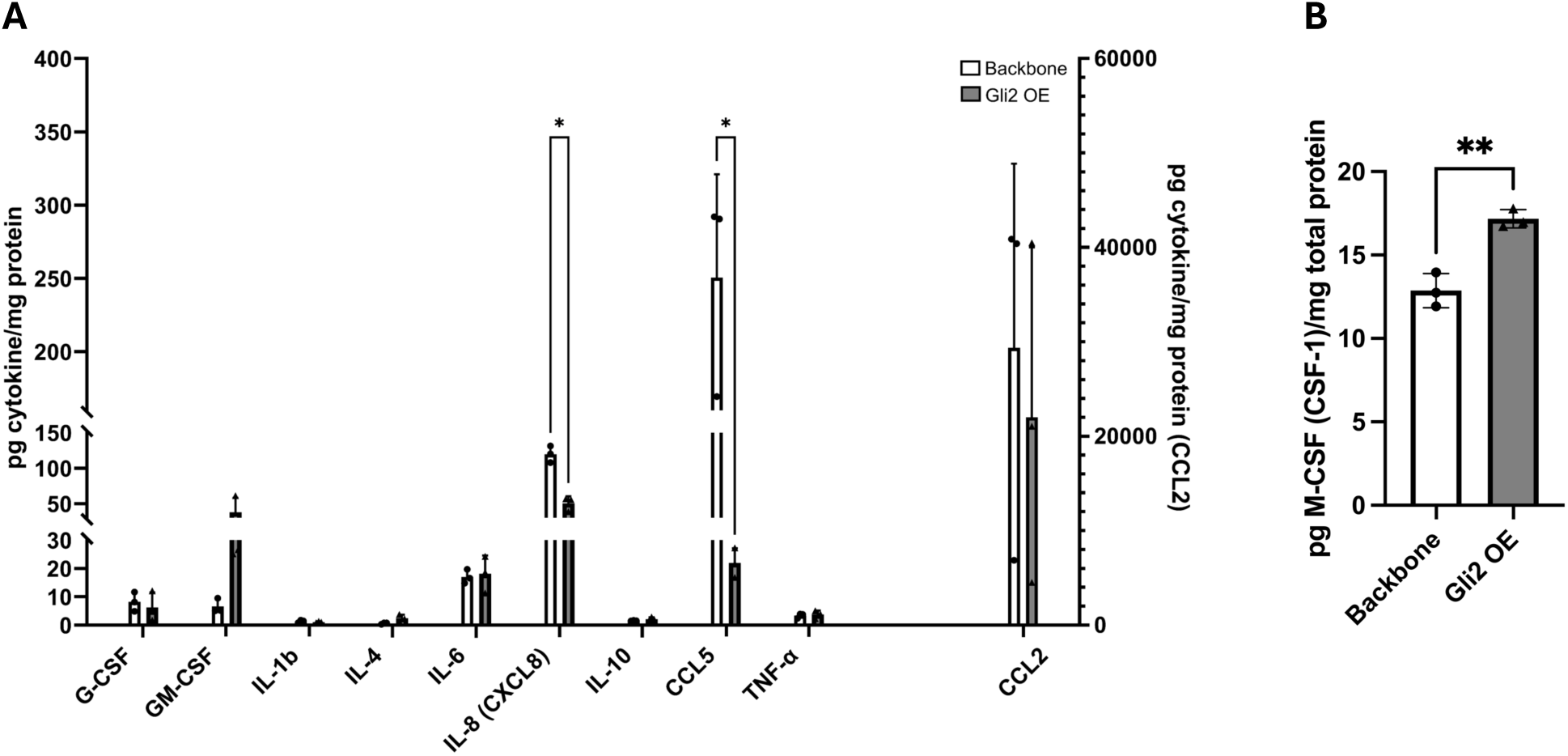
Gli2 overexpression results in a cytokine secretion profile distinct from backbone SW872 cells. (A) MILLIPLEX MAP Human Cytokine/Chemokine Magnetic Bead Panel results showing cytokine concentrations in conditioned media from SW872 backbone and Gli2 OE cells. (B) ELISA for M-CSF (CSF-1) was performed on the same conditioned media. Mean and standard deviation shown. *q < 0.05. **p < 0.01. Biological replicates from three independent experiments.

### Orthotopic SW872 Gli2 OE tumors have an altered gross phenotype

SW872 backbone and Gli2 OE cells were then bilaterally injected into the dorsal/posterior aspect of the inguinal fat pads of nude mice and allowed to establish for 18 days (**Figure 6A**). Over the course of these 18 days, there was no significant difference in mouse body weight (**Figure 6B**). Upon sacrifice, the entirety of both fat pads was excised. There was no significant difference in fat pad weight (**Figure 6C**). However, the gross phenotype of the tumor-bearing fat pad types was altered, with Gli2 OE tumor-bearing fat pads appearing redder in color, potentially suggesting increased vascularization (**Figure 6D**).

**Figure 6.**
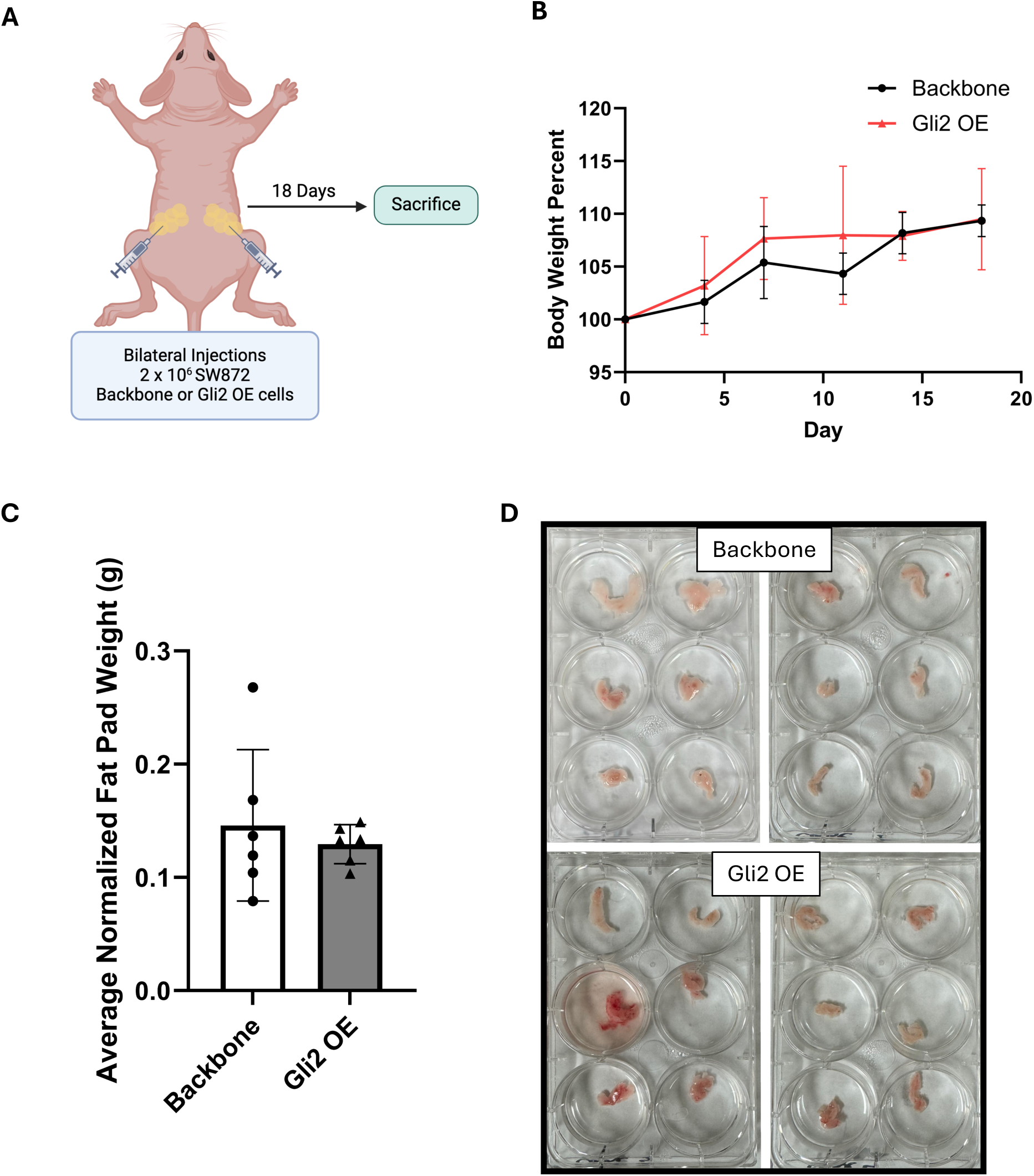
Orthotopic dedifferentiated liposarcoma tumors with overexpressed Gli2 do not increase fat pad weight but have an altered gross phenotype. (A) Schematic of mouse study. Gli2 overexpressing SW872 cells or backbone control cells were injected bilaterally into the dorsal/posterior aspect of the inguinal fat pad. Fat pads were collected at sacrifice 18 days later. (B) Mouse weight over time as percent of original weight. (C) Final weight of tumor-bearing fat pad after sacrifice normalized to mouse weight. (D) Image of resected fat pads (top left = backbone male, top right = backbone female, bottom left = Gli2 OE male, bottom right = Gil2 OE female). Mean and standard deviation shown. n = 6. Schematic created with BioRender.com

### Gli2 OE tumor-bearing fat pads contain macrophages with an immunosuppressive phenotype

The SW872 tumor-bearing fat pads were then processed via enzymatic digestion and evaluated for murine myeloid surface markers by flow cytometry. There was no significant difference in CD11b expression among CD45+ lymphocytes (**Figure 7A**) or F4/80+ macrophages (**Figure 7B**) between the two tumor groups. However, among F4/80+ macrophages present in the tumors, there were significant phenotypic differences. Gli2 OE tumor-bearing fat pads had significantly higher levels of CD163+ macrophages (**Figure 7C**) and CD206+ macrophages (**Figure 7D**), suggesting an increase in immunosuppressive M2-like macrophages. Interestingly, CCR2+ macrophages also increased in the Gli2 OE group, although comprising a smaller percentage of total macrophages than the CD163+ or CD206+ macrophages (**Figure 7E**).

**Figure 7.**
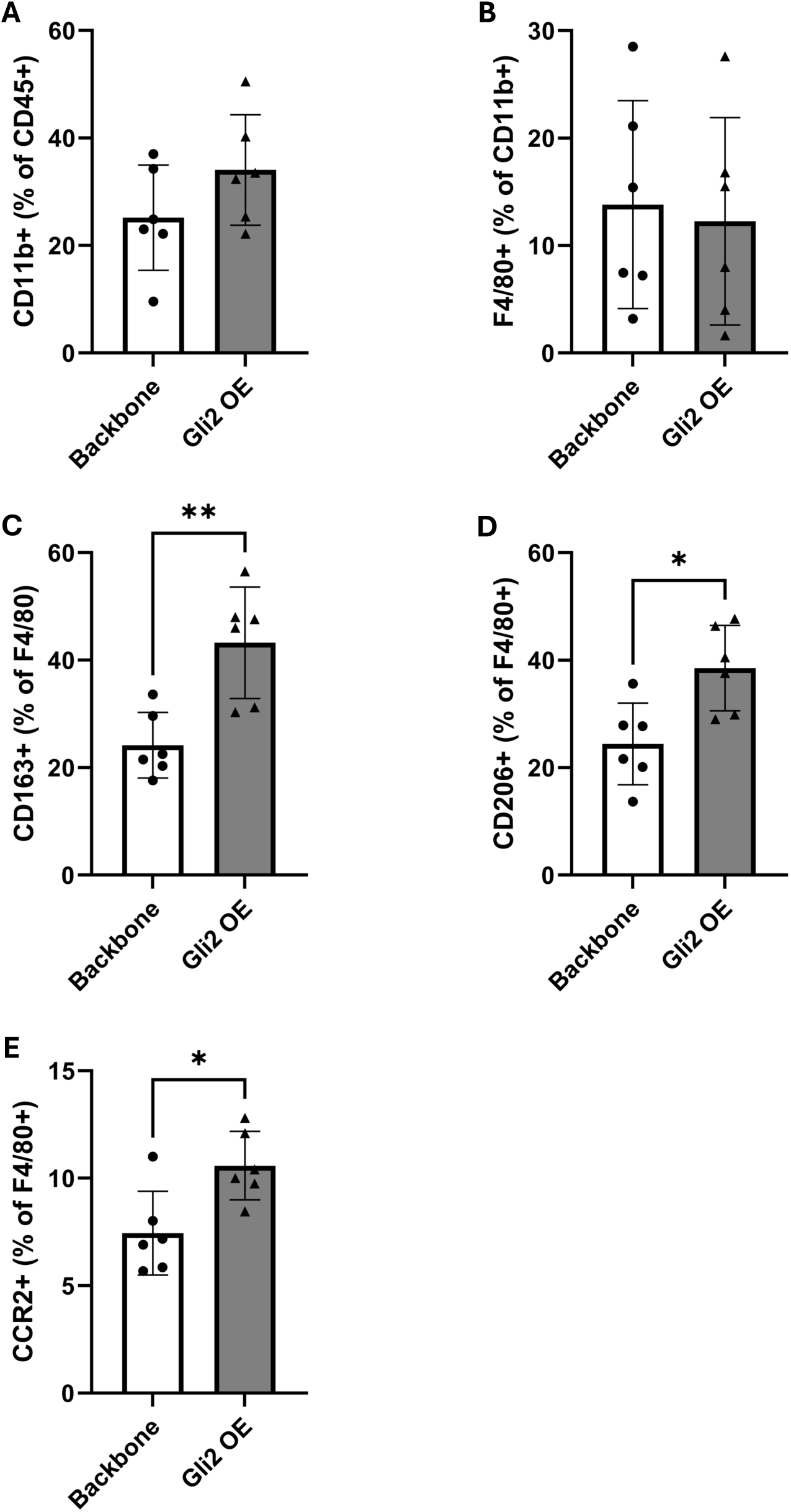
Fat pads with Gli2 OE tumors have an altered macrophage phenotype. Dorsal fat pads with SW872 backbone or Gli2 OE tumors were collected 18 days after tumor injection, and flow cytometry was performed for mouse myeloid phenotypic markers. (A) Percent of CD11b+ cells within CD45+ population. (B) Percent of F4/80+ cells within CD11b+ population. (C) Percent of CD163+ cells within F4/80+ population. (D) Percent of CD206+ cells within F4/80+ population. (E) Percent of CCR2+ cells within F4/80+ population. Mean and standard deviation shown. *p < 0.05. **p < 0.01. n = 6

Although CCR2 is sometimes considered a marker of M1-like macrophages, myeloid cells expressing CCR2 in the tumor microenvironment often exhibit immunosuppressive properties upon recruitment by its tumor-secreted ligand CCL2[33], which is abundantly expressed by both SW872 backbone and Gli2 OE cells as shown in **Figure 5A**.

Finally, immature myeloid cells were evaluated via expression of Ly6C and Ly6G among CD11b+ cells (**Figure S4**). There was no significant difference in monocytic Ly6C+/Ly6G-cells between groups, although CD163, CD206, and CCR2 positivity followed similar trends as those seen in mature F4/80+ macrophages. Granulocytic immature myeloid cells (Ly6C-/Ly6G+) were also evaluated but comprised less than 1% of CD11b cells in each sample. In total, the above data suggests that Gli2 OE tumor-bearing fat pads are home to macrophages with an increasingly immunosuppressive phenotype compared to those with SW872 backbone tumors.

## Discussion and Conclusions

Dedifferentiated liposarcomas are clinically challenging neoplasms that exemplify the role of differentiation status in affecting tumor aggressiveness. Our previous work has demonstrated that DDLPS tumors have increased Hedgehog signaling and have overlapping gene signatures with adipose progenitor populations[7]. This work also demonstrated a notable inverse correlation between Gli2 expression and HLA expression, T-cell markers, and myeloid markers. Therefore, in this work, we sought to explore potential mechanisms by which this phenotype may occur, whether due to Gli2 induction of a stem-like phenotype with low HLA expression or differentiation-extrinsic mechanisms such as altered cytokine expression.

We established a Gli2-overexpressing SW872 DDLPS cell line and proceeded with phenotypic characterization of these cells. Although Gli2 OE cells had an increased proliferative capacity and reduced adipogenic potential, they had a decreased self-renewal capacity, which would be unexpected if Gli2 was truly driving an MSC-like phenotype in these cells (**Figure 1**). Bulk RNA sequencing further characterized the distinct transcriptional profile that resulted from Gli2 overexpression (**Figure 2**). Of particular interest was enrichment in inflammatory pathways and processes related to skeletal development. The latter especially prompted us to proceed with a more detailed evaluation of the differentiation state of these cells as shown in **Figure 3**. Neither cell line expressed genes for the adipocyte markers adiponectin or leptin, and there was a nonsignificant decrease in the expression of lipoprotein lipase in Gli2 OE cells.

Interestingly, Gli2 OE cells had decreased levels of several mesenchymal stem cell lineage markers. Because of the role of Gli2 in MSC differentiation toward osteoblasts[16,21–23], we then evaluated markers of this lineage, which demonstrated an increase in osteopontin and sclerostin. Although sclerostin, a marker of osteocytes, has been shown to increase adipocyte differentiation[24], literature suggests that in skeletal sarcomas, it is positively correlated with osteogenesis[25]. Furthermore, Gli2 OE cells had increased expression of *NOTCH3*, *HEY1*, and *JAG1* (**Figure S3**), all components of the Notch signaling pathway which increases osteoblast differentiation and decreases adipogenesis[30,31]. In total, this data demonstrates that rather than driving DDLPS cells toward an MSC-like phenotype, Gli2 overexpression appears to move the cells past an MSC-like state and toward osteoblastic differentiation. This observation may be explored in future studies as a mechanism for the ossification sometimes observed in patient DDLPS tumors[34].

We then evaluated the expression of antigen presentation genes and cytokines in these DDLPS cells. While HLA class I expression, particularly HLA-C, appeared to decrease in Gli2 OE cells, more notable was the differential expression of several cytokines, especially *CSF1* and *CSF2*, which significantly increased in Gli2 OE cells (**Figure 4**). Secreted cytokines were also altered by Gli2 overexpression, specifically a decrease in IL-8 and CCL5 and an increase in M-CSF (**Figure 5**). Although IL-8 and CCL5 have pleiotropic roles in shaping the tumor-immune microenvironment[35,36], M-CSF is notable for recruiting monocytic populations and inducing M2-like polarization of tumor-associated macrophages[37]. Given the evident role of Gli2 overexpression in cytokine expression as well as tumor cell differentiation status, we then proceeded with *in vivo* evaluation of the tumor microenvironment.

We established an orthotopic model of dedifferentiated liposarcoma by injecting SW872 backbone and Gli2 OE cells into the dorsal fat pads of nude mice. After 18 days of tumor establishment, the fat pads were collected for downstream analysis. Grossly, Gli2 OE tumor-bearing fat pads were the same weight as SW872 backbone tumors but had a qualitatively altered appearance (**Figure 6**). Because of the changes in cytokine expression and secretion *in vitro*, we anticipated changes in the phenotype of tumor-associated myeloid cells (**Figure 7**). Indeed, there was a significant increase in CD163 and CD206 positive macrophages in the Gli2 OE tumor-bearing fat pad by flow cytometry, suggesting M2-like macrophage polarization. This may bear relevance to patient tumors, as dedifferentiated components of DDLPS tumors harbor more M2-like macrophages than well-differentiated components of the same tumors[20]. There was also a significant increase in CCR2+ macrophages in the Gli2 OE group, which were likely recruited by the notably high levels of CCL2 secreted by tumor cells as shown in **Figure 5**. CCL2 has been shown recruit macrophages to adipose tissue[38,39], so a similar mechanism may be occurring in this DDLPS model.

In summary, this study demonstrates that Gli2 overexpression drives SW872 DDLPS cells toward a genetic signature indicative of osteoblast differentiation with a decrease in mesenchymal markers. These cells also have a distinct transcriptional and secretory cytokine profile, and establishment and growth of SW872 Gli2 OE cells in an orthotopic model of DDLPS results in increased recruitment or polarization of M2-like macrophages. The true mechanism by which Gli2 associates with an immunosuppressive phenotype in DDLPS tumors may be a combination of these factors, as tumor cell phenotype, functional cytokine secretion, and resultant tumor immunophenotype are likely inextricably linked.

These findings present several potential routes of future investigation. First, *in vivo* validation of the *in vitro* findings of osteoblastic lineage marker enhancement in Gli2 OE cells can be explored through histological analysis of tumor-bearing fat pads, including staining for mineralization in the fat pads at a longer timepoint after tumor inoculation. Additionally, incorporation of CCL2 or CSF-1 inhibitory antibodies *in vivo* could better elucidate the role of individual cytokines in recruiting immunosuppressive macrophages to the DDLPS tumor. Further mechanistic studies will be important to stratify the relative importance of tumor cell differentiation state versus cytokine secretion in inducing the immunosuppressive myeloid populations observed. There are additional roles for emerging therapies in future studies. For example, CD47, the “don’t eat me” signal that is expressed in most soft tissue sarcoma and interacts with SIRPα on myeloid cells to inhibit phagocytosis[40], is another promising target to inhibit Gli2-related immunosuppression. A recent study using cell suspensions from patient soft tissue sarcoma cells demonstrated that anti-CD47 therapy induced pro-inflammatory cytokine secretion[40]. Finally, mechanistic evaluation of T-cell response to tumor cell Gli2 overexpression is challenging, as there are no readily available mouse DDLPS cell lines to use in an immunocompetent mouse model. However, future studies utilizing patient tumors may identify additional correlations between Gli2 expression and immune infiltration.

## Author Contributions (CRediT)

NEB: Conceptualization, Methodology, Validation, Investigation, Writing – Original Draft, Visualization; EPB: Conceptualization, Methodology, Validation, Investigation, Writing – Original Draft, Visualization; DVP: Methodology, Investigation, Writing – Review & Editing; EJC: Methodology, Investigation, Writing – Review & Editing; MAC: Methodology, Investigation, Writing – Review & Editing; JEB: Investigation, Writing – Review & Editing; JSM: Investigation, Writing – Review & Editing; JAS: Investigation, Writing – Review & Editing; JAR: Conceptualization, Resources, Writing – Review & Editing, Supervision, Funding Acquisition

## Acknowledgements

RNA sequencing was completed by the Vanderbilt Technologies for Advanced Genomics (VANTAGE) core laboratory for next-generation sequencing, which is supported in part by Clinical and Translational Science Award Grant 5UL1 RR024975-03, Vanderbilt Ingram Cancer Center Grant P30 CA68485, Vanderbilt Vision Center Grant P30 EY08126, and National Institutes of Health/National Center for Research Resources Grant G20 RR030956. Luminex assays were performed by the VUMC Hormone Assay and Analytical Services Core which is supported by NIH grants DK059637 and DK020593. The authors also thank Dr. Christian Warren in the flow cytometry core at the United States Department of Veterans Affairs, Tennessee Valley Healthcare System, Nashville, TN for his assistance with flow cytometry experiments. The authors also thank Dr. Dixon Dorand (Dr. Ben Park lab) for reagents and technical advice.

This work was supported by the Veterans Health Administration under grant number 5I01BX001957 and the National Institutes of Health under award numbers 1R01CA264508 and T32GM007347. The content is solely the responsibility of the authors and does not necessarily represent the official views of the National Institutes of Health.

## SUPPLEMENTARY TABLES AND FIGURES

**Table S1.** Taqman primers used for qRT-PCR in SW872 *in vitro* studies.

| Assay ID | Gene | Assay ID | Gene | Assay ID | Gene |
| --- | --- | --- | --- | --- | --- |
| Hs99999901_s1 | 18S | Hs00266645_m1 | FGF2 | Hs00236988_g1 | MIF |
| Hs00605917_m1 | ADIPOQ | Hs00270951_s1 | FOXC2 | Hs00707120_s1 | NES |
| Hs01029144_m1 | ALPL | Hs01054576_m1 | FOXO1 | Hs01062014_m1 | NOTCH1 |
| Hs00187842_m1 | B2M | Hs01110766_m1 | GLI1 | Hs01050702_m1 | NOTCH2 |
| Hs00237013_m1 | CCL11 | Hs01119974_m1 | GLI2 | Hs01128541_m1 | NOTCH3 |
| Hs00234140_m1 | CCL2 | Hs00172878_m1 | HES1 | Hs00965889_m1 | NOTCH4 |
| Hs00355476_m1 | CCL20 | Hs00232618_m1 | HEY1 | Hs00159686_m1 | NT5E |
| Hs99999148_m1 | CCL4 | Hs01011015_m1 | HHIP | Hs00931793_m1 | OSX |
| Hs00982282_m1 | CCL5 | Hs01058806_g1 | HLA-A | Hs00966522_m1 | PDGFB |
| Hs00171147_m1 | CCL7 | Hs00818803_g1 | HLA-B | Hs01115513_m1 | PPARG |
| Hs02379687_s1 | CD24 | Hs00740298_g1 | HLA-C | Hs00181117_m1 | PTCH1 |
| Hs00204257_m1 | CD274 | Hs00256884_m1 | HOXB4 | Hs01047973_m1 | RUNX2 |
| Hs01567185_m1 | CD36 | Hs00989291_m1 | IFNG | Hs01090242_m1 | SMO |
| Hs01075864_m1 | CD44 | Hs00961622_m1 | IL10 | Hs00195591_m1 | SNAI1 |
| Hs01023894_m1 | CDH1 | Hs00174148_m1 | IL11 | Hs00950344_m1 | SNAI2 |
| Hs00983056_m1 | CDH2 | Hs01073447_m1 | IL12A | Hs00228830_m1 | SOST |
| Hs00270923_s1 | CEBPB | Hs01011518_m1 | IL12B | Hs00959010_m1 | SPP1 |
| Hs00174164_m1 | CSF1 | Hs00174379_m1 | IL13 | Hs00388675_m1 | TAP1 |
| Hs00929873_m1 | CSF2 | Hs00174383_m1 | IL17A | Hs00998133_m1 | TGFB1 |
| Hs00738432_g1 | CSF3 | Hs01038788_m1 | IL18 | Hs00234244_m1 | TGFB2 |
| Hs00236937_m1 | CXCL1 | Hs00174092_m1 | IL1A | Hs01086000_m1 | TGFB3 |
| Hs00171042_m1 | CXCL10 | Hs01555410_m1 | IL1B | Hs00174816_m1 | THY1 |
| Hs00171138_m1 | CXCL11 | Hs00174114_m1 | IL2 | Hs00174128_m1 | TNF |
| Hs03676656_mH | CXCL12 | Hs00372324_m1 | IL23A | Hs00361186_m1 | TWIST1 |
| Hs00601975_m1 | CXCL2 | Hs00174122_m1 | IL4 | Hs00263977_m1 | WNT3A |
| Hs01099660_g1 | CXCL5 | Hs00174131_m1 | IL6 | Hs00362452_m1 | WNT6 |
| Hs00174103_m1 | CXCL8 | Hs01070032_m1 | JAG1 | Hs00232783_m1 | ZEB1 |
| Hs00171065_m1 | CXCL9 | Hs00174877_m1 | LEP | Hs00207691_m1 | ZEB2 |
| Hs01009390_m1 | DMP1 | Hs00173425_m1 | LPL |  |  |

**Table S2.** Flow cytometry antibodies and dilutions for SW872 *in vivo* study.

| Target | Fluorophore | Clone | Supplier/Catalog Number | Dilution |
| --- | --- | --- | --- | --- |
| Anti-mouse CD45 | APC-Cy7 | 30-F11 | BD 557659 | 1:1000 |
| Anti-mouse CD11b | FITC | M1/70 | Biolegend 101206 | 1:2500 |
| Anti-mouse F4/80 | AF700 | BM8 | Biolegend 123129 | 1:500 |
| Anti-mouse Gr1/Ly6G | PerCP-Cy5.5 | 1A8 | Biolegend 127615 | 1:500 |
| Anti-mouse Ly6C | BV605 | HK1.4 | Biolegend 128036 | 1:200 |
| Anti-mouse CD192<br>(CCR2) | BV421 | SA203G11 | Biolegend 150605 | 1:100 |
| Anti-mouse CD206 | PE | C068C2 | Biolegend 141705 | 1:250 |
| Anti-mouse CD163 | PE-Cy7 | S15049F | Biolegend 156707 | 1:100 |

**Figure S1.**
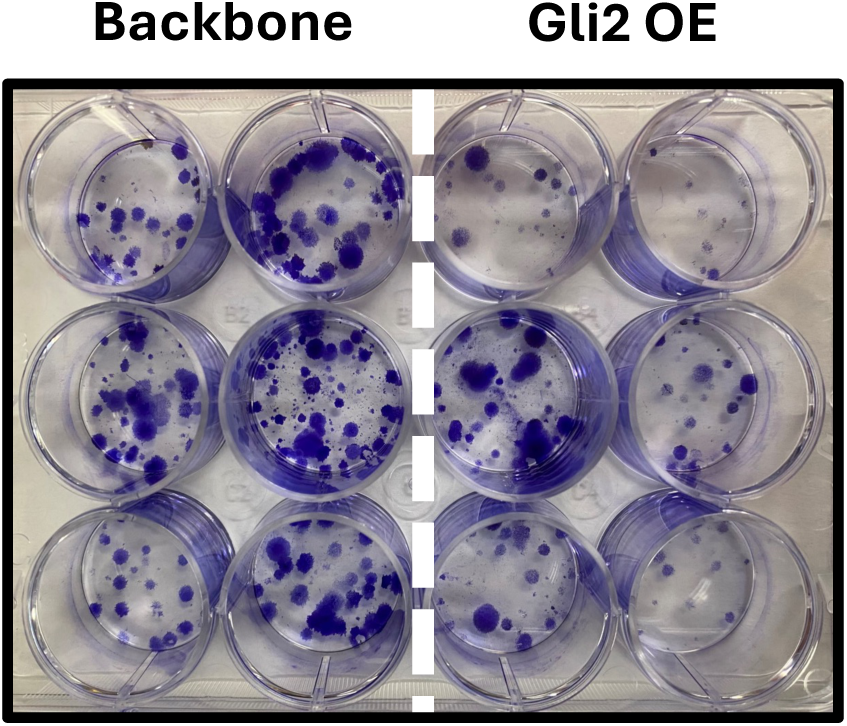
Colony forming assay demonstrates that SW872 Gli2 OE cells have reduced capacity for self-renewal compared to backbone control cells. Cells were plated at a low density and cultured for 14 days followed by staining with crystal violet. The above is a representative image of the data plotted in Figure 1D.

**Figure S2.**
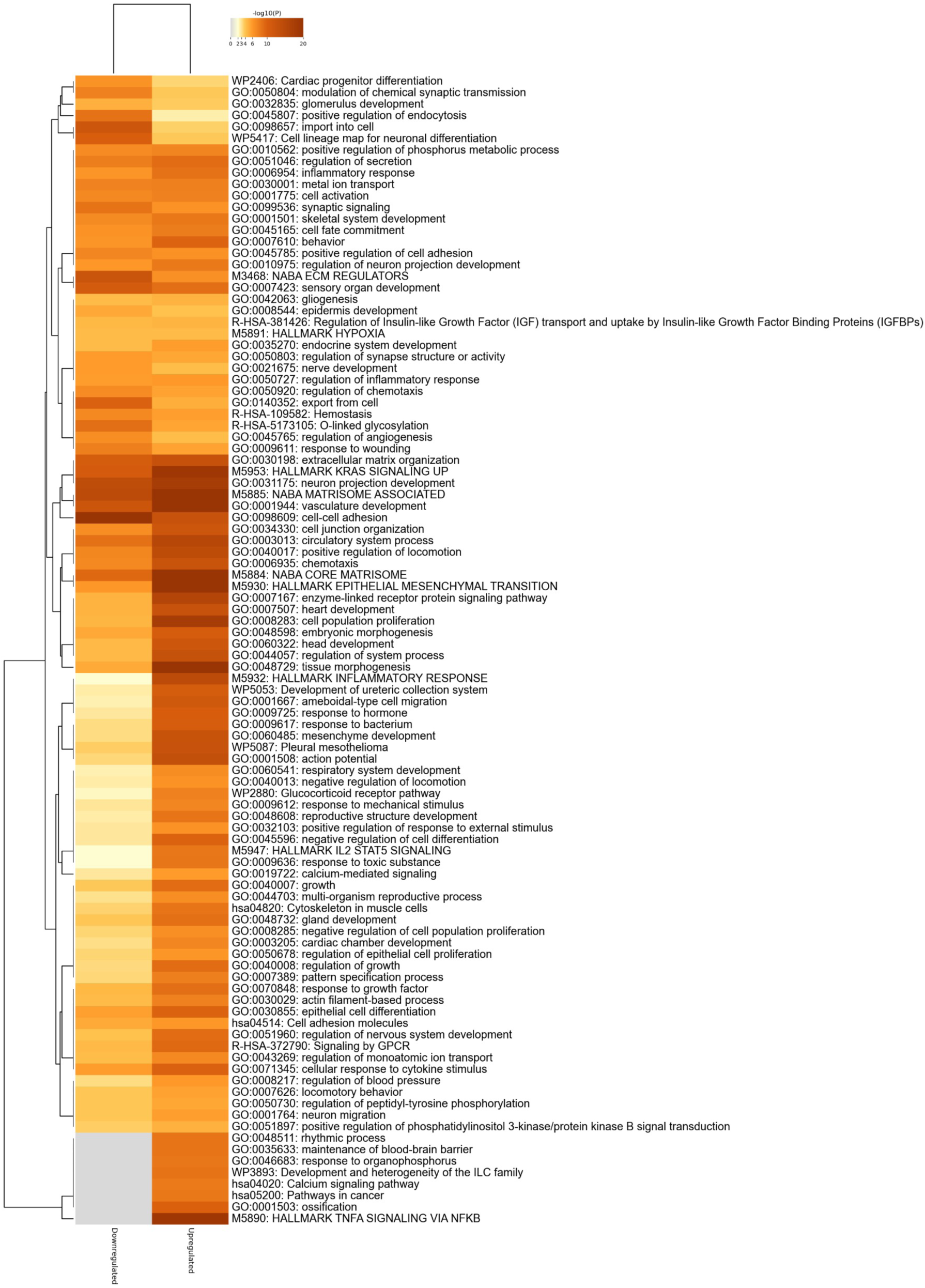
Expanded heatmap of the top 100 enriched clusters for downregulated and upregulated genes in SW872 Gli2 OE cells vs SW872 control cells. SW872 cells were transfected with either Gli2 overexpression vector or empty vector (backbone) control. Bulk RNA sequencing was performed, identifying 937 unique downregulated genes and 1304 unique upregulated genes. Metascape was used to perform Gene Ontology and pathway enrichment analyses, and the top 100 enriched clusters for downregulated and upregulated genes are shown.

**Figure S3.**
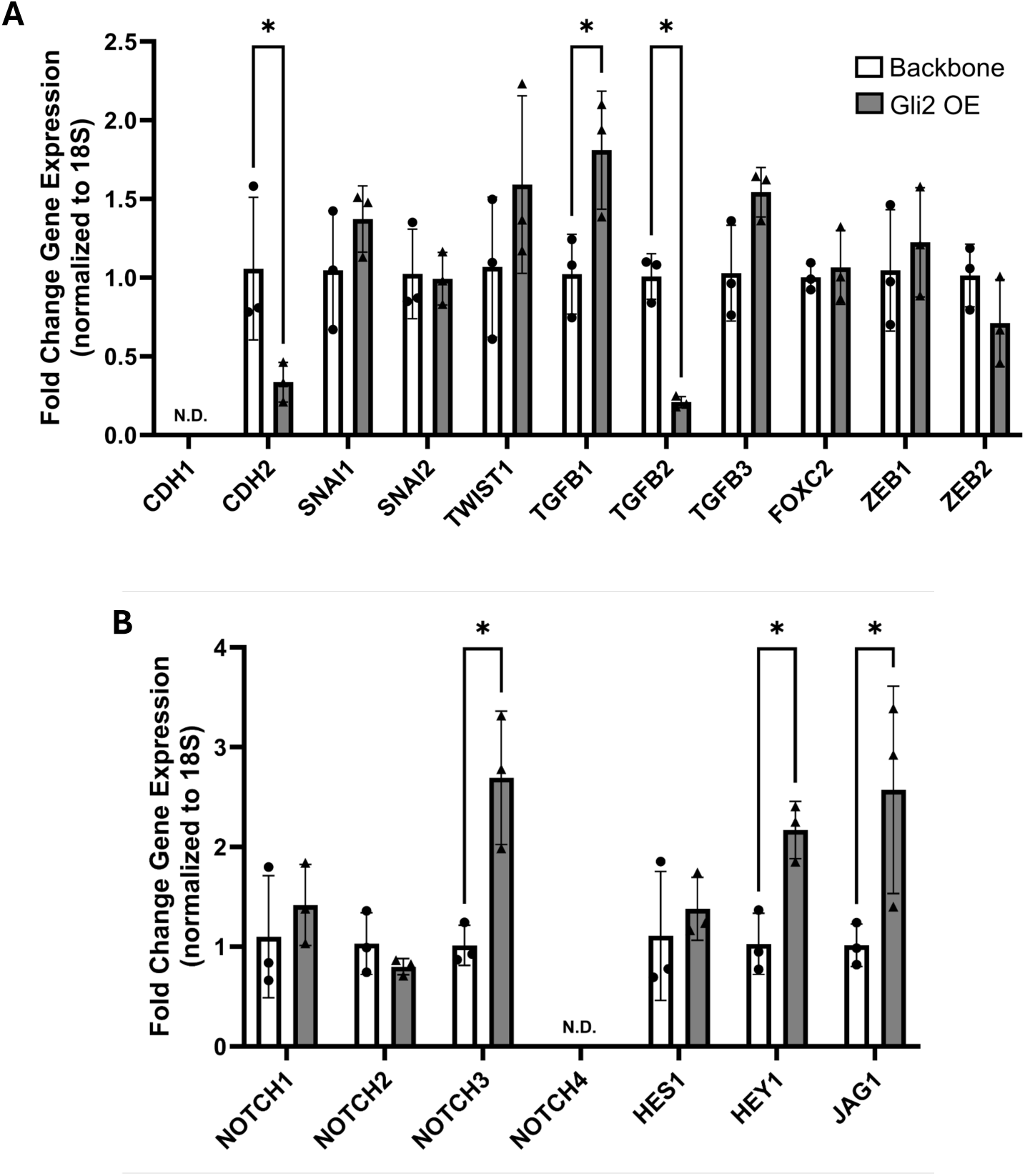
Gli2 overexpressing SW872 cells have reduced mesenchymal marker expression and increased Notch signaling marker expression. Gene expression of (A) epithelial-mesenchymal transition genes and (B) Notch signaling genes. *q < 0.05. Biological replicates from three independent experiments. N.D. = not detected.

**Figure S4.**
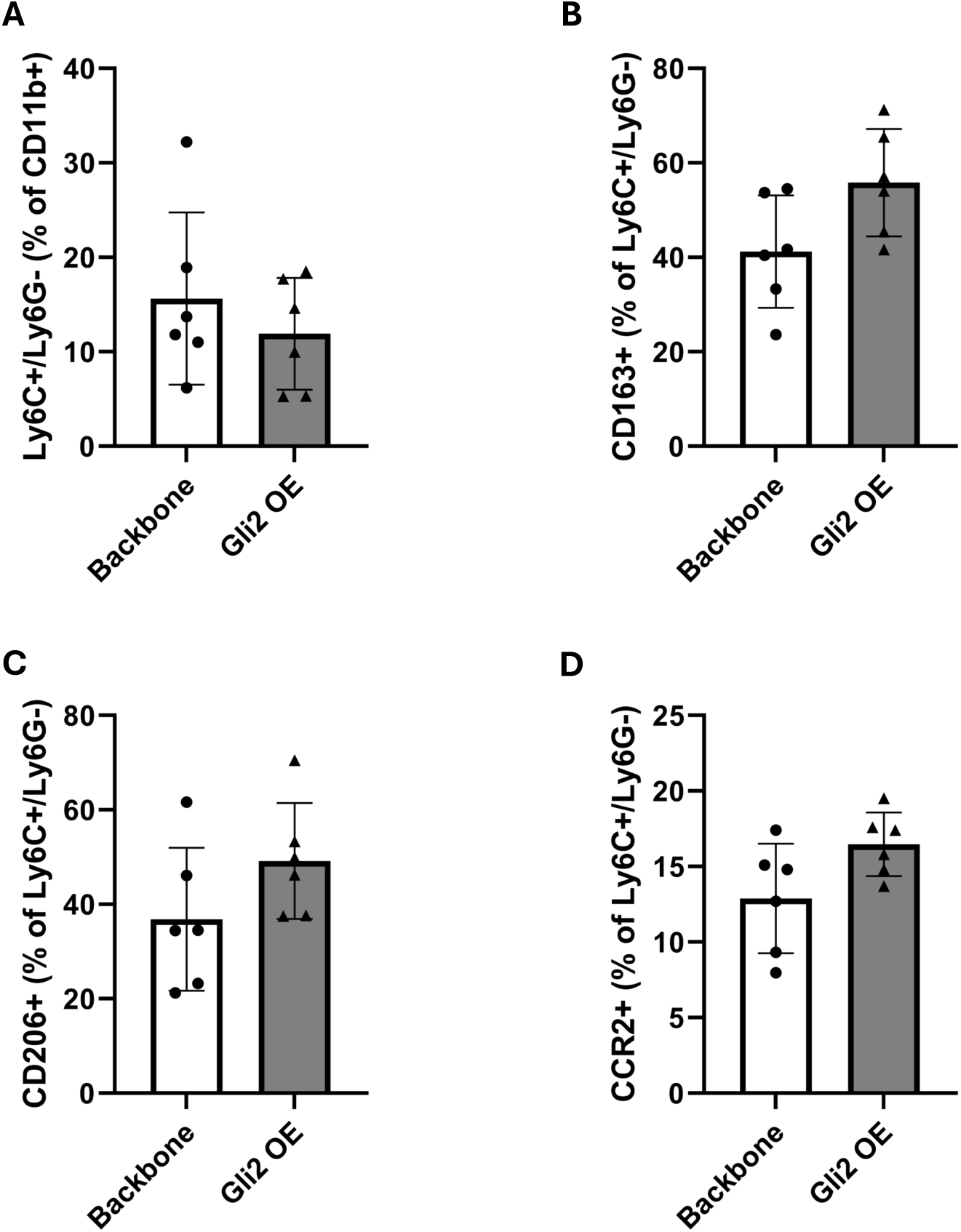
SW872 backbone and Gli2 OE cells have similar populations of immature myeloid cells. (A) Percent of Ly6C+/Ly6G-cells within CD11b+ population. (B-D) Percent of CD163+, CD206+, and CCR2+ cells within CD11b+/ Ly6C+/Ly6G-population. n = 6

## REFERENCES

1. Damerell V, Pepper MS, Prince S. Molecular mechanisms underpinning sarcomas and implications for current and future therapy. Signal Transduct Target Ther. 2021;6(1):1–19. doi:10.1038/s41392-021-00647-8

2. Thway K. Well-differentiated liposarcoma and dedifferentiated liposarcoma: An updated review. Semin Diagn Pathol. 2019;36(2):112–121. doi:10.1053/J.SEMDP.2019.02.006

3. Jones RL, Lee ATJ, Thway K, Huang PH. Clinical and molecular spectrum of liposarcoma. Journal of Clinical Oncology. 2018;36(2):151–159. doi:10.1200/JCO.2017.74.959

4. Singer S, Antonescu CR, Riedel E, Brennan MF, Pollock RE. Histologic Subtype and Margin of Resection Predict Pattern of Recurrence and Survival for Retroperitoneal Liposarcoma. Ann Surg. 2003;238(3):358. doi:10.1097/01.SLA.0000086542.11899.38

5. Han D, Rodriguez-Bravo V, Charytonowicz E, et al. Targeting Sarcoma Tumor-Initiating Cells through Differentiation Therapy. Stem Cell Res. 2017;21:117. doi:10.1016/J.SCR.2017.04.004

6. Wang Y, Huang J, Gong L, et al. The Plasticity of Mesenchymal Stem Cells in Regulating Surface HLA-I. iScience. 2019;15:66–78. doi:10.1016/J.ISCI.2019.04.011

7. Beadle EP, Bennett NE, Rhoades JA. Bioinformatics Screen Reveals Gli-Mediated Hedgehog Signaling as an Associated Pathway to Poor Immune Infiltration of Dedifferentiated Liposarcoma. Cancers (Basel). 2023;15(13):3360. doi:10.3390/cancers15133360

8. Ewels PA, PelÅer A, Fillinger S, et al. The nf-core framework for community-curated bioinformatics pipelines. Nature Biotechnology 2020 38:3. 2020;38(3):276-278. doi:10.1038/s41587-020-0439-x

9. Love MI, Anders S, Kim V, Huber W. RNA-Seq workflow: gene-level exploratory analysis and differential expression. F1000Res. 2015;4:1070. doi:10.12688/F1000RESEARCH.7035.1

10. Love MI, Huber W, Anders S. Moderated estimation of fold change and dispersion for RNA-seq data with DESeq2. Genome Biol. 2014;15(12):1–21. doi:10.1186/s13059-014-0550-8

11. Zhou Y, Zhou B, Pache L, et al. Metascape provides a biologist-oriented resource for the analysis of systems-level datasets. Nature Communications 2019 10:1. 2019;10(1):1-10. doi:10.1038/s41467-019-09234-6

12. Stratford EW, Castro R, Daffinrud J, et al. Characterization of Liposarcoma Cell Lines for Preclinical and Biological Studies. Sarcoma. 2012;2012. doi:10.1155/2012/148614

13. Cui J, Dean D, Hornicek FJ, Pollock RE, Hoffman RM, Duan Z. ATR inhibition sensitizes liposarcoma to doxorubicin by increasing DNA damage. Am J Cancer Res. 2022;12(4):1577. /pmc/articles/PMC9077062/?report=abstract

14. Ikram MS, Neill GW, Regl G, et al. GLI2 is expressed in normal human epidermis and BCC and induces GLI1 expression by binding to its promoter. Journal of Investigative Dermatology. 2004;122(6):1503–1509. doi:10.1111/j.0022-202X.2004.22612.x

15. Sigafoos AN, Paradise BD, Fernandez-Zapico ME. Hedgehog/GLI Signaling Pathway: Transduction, Regulation, and Implications for Disease. Cancers (Basel). 2021;13(14). doi:10.3390/CANCERS13143410

16. Spinella-Jaegle S, Rawadi G, Kawai S, et al. Sonic hedgehog increases the commitment of pluripotent mesenchymal cells into the osteoblastic lineage and abolishes adipocytic differentiation. J Cell Sci. 2001;114(11):2085–2094. doi:10.1242/JCS.114.11.2085

17. Zhang L, Fu X, Ni L, et al. Hedgehog Signaling Controls Bone Homeostasis by Regulating Osteogenic/Adipogenic Fate of Skeletal Stem/Progenitor Cells in Mice. Journal of Bone and Mineral Research. 2022;37(3):559–576. doi:10.1002/JBMR.4485

18. Fontaine C, Cousin W, Plaisant M, Dani C, Peraldi P. Hedgehog Signaling Alters Adipocyte Maturation of Human Mesenchymal Stem Cells. Stem Cells. 2008;26(4):1037–1046. doi:10.1634/STEMCELLS.2007-0974

19. Varjosalo M, Taipale J. Hedgehog: functions and mechanisms. Genes Dev. 2008;22(18):2454-2472. doi:10.1101/GAD.1693608

20. Gruel N, Quignot C, Lesage L, et al. Cellular origin and clonal evolution of human dedifferentiated liposarcoma. Nature Communications 2024 15:1. 2024;15(1):1-18. doi:10.1038/s41467-024-52067-1

21. Zhao M, Qiao M, Harris SE, Chen D, Oyajobi BO, Mundy GR. The Zinc Finger Transcription Factor Gli2 Mediates Bone Morphogenetic Protein 2 Expression in Osteoblasts in Response to Hedgehog Signaling. Mol Cell Biol. 2006;26(16):6197. doi:10.1128/MCB.02214-05

22. Xu L, Ji C, Yu T, Luo J. The effects of Gli1 and Gli2 on BMP9-induced osteogenic differentiation of mesenchymal stem cells. Tissue Cell. 2023;84:102168. doi:10.1016/J.TICE.2023.102168

23. Shimoyama A, Wada M, Ikeda F, et al. Ihh/Gli2 Signaling Promotes Osteoblast Differentiation by Regulating Runx2 Expression and Function. Mol Biol Cell. 2007;18(7):2411. doi:10.1091/MBC.E06-08-0743

24. Ukita M, Yamaguchi T, Ohata N, Tamura M. Sclerostin Enhances Adipocyte Differentiation in 3T3-L1 Cells. J Cell Biochem. 2016;117(6):1419–1428. doi:10.1002/JCB.25432

25. Shen J, Meyers CA, Shrestha S, et al. Sclerostin expression in skeletal sarcomas. Hum Pathol. 2016;58:24. doi:10.1016/J.HUMPATH.2016.07.016

26. Xu L, Meng F, Ni M, Lee WYW, Li G. N-cadherin regulates osteogenesis and migration of bone marrow-derived mesenchymal stem cells. Mol Biol Rep. 2013;40(3):2533–2539. doi:10.1007/S11033-012-2334-0

27. Wu M, Wu S, Chen W, Li YP. The roles and regulatory mechanisms of TGF-β and BMP signaling in bone and cartilage development, homeostasis and disease. Cell Research 2024 34:2. 2024;34(2):101-123. doi:10.1038/s41422-023-00918-9

28. Kasagi S, Chen W. TGF-beta1 on osteoimmunology and the bone component cells. Cell Biosci. 2013;3(1):1–7. doi:10.1186/2045-3701-3-4

29. Spinella-Jaegle S, Roman-Roman S, Faucheu C, et al. Opposite effects of bone morphogenetic protein-2 and transforming growth factor-β1 on osteoblast differentiation. Bone. 2001;29(4):323–330. doi:10.1016/S8756-3282(01)00580-4

30. Song BQ, Chi Y, Li X, et al. Inhibition of Notch Signaling Promotes the Adipogenic Differentiation of Mesenchymal Stem Cells Through Autophagy Activation and PTEN-PI3K/AKT/mTOR Pathway. Cellular Physiology and Biochemistry. 2015;36(5):1991–2002. doi:10.1159/000430167

31. Ugarte F, Ryser M, Thieme S, et al. Notch signaling enhances osteogenic differentiation while inhibiting adipogenesis in primary human bone marrow stromal cells. Exp Hematol. 2009;37(7):867–875.e1. doi:10.1016/j.exphem.2009.03.007

32. Dahlman I, Kaaman M, Olsson T, et al. A Unique Role of Monocyte Chemoattractant Protein 1 among Chemokines in Adipose Tissue of Obese Subjects. J Clin Endocrinol Metab. 2005;90(10):5834–5840. doi:10.1210/JC.2005-0369

33. Fei L, Ren X, Yu H, Zhan Y. Targeting the CCL2/CCR2 Axis in Cancer Immunotherapy: One Stone, Three Birds? Front Immunol. 2021;12:771210. doi:10.3389/fimmu.2021.771210

34. Yamashita K, Kohashi K, Yamada Y, et al. Osteogenic differentiation in dedifferentiated liposarcoma: a study of 36 cases in comparison to the cases without ossification. Histopathology. 2018;72(5):729–738. doi:10.1111/HIS.13421

35. Aldinucci D, Borghese C, Casagrande N. The CCL5/CCR5 Axis in Cancer Progression. Cancers (Basel). 2020;12(7):1–30. doi:10.3390/CANCERS12071765

36. Fousek K, Horn LA, Palena C. Interleukin-8: a Chemokine at the Intersection of Cancer Plasticity, Angiogenesis, and Immune Suppression. Pharmacol Ther. 2021;219:107692. doi:10.1016/J.PHARMTHERA.2020.107692

37. Hume DA, MacDonald KPA. Therapeutic applications of macrophage colony-stimulating factor-1 (CSF-1) and antagonists of CSF-1 receptor (CSF-1R) signaling. Blood. 2012;119(8):1810–1820. doi:10.1182/BLOOD-2011-09-379214

38. Kamei N, Tobe K, Suzuki R, et al. Overexpression of monocyte chemoattractant protein-1 in adipose tissues causes macrophage recruitment and insulin resistance. Journal of Biological Chemistry. 2006;281(36):26602–26614. doi:10.1074/jbc.M601284200

39. Kanda H, Tateya S, Tamori Y, et al. MCP-1 contributes to macrophage infiltration into adipose tissue, insulin resistance, and hepatic steatosis in obesity. Journal of Clinical Investigation. 2006;116(6):1494. doi:10.1172/JCI26498

40. Ozaniak A, Smetanova J, Bartolini R, et al. A novel anti-CD47-targeted blockade promotes immune activation in human soft tissue sarcoma but does not potentiate anti-PD-1 blockade. J Cancer Res Clin Oncol. 2023;149(7):3789–3801. doi:10.1007/s00432-022-04292-8

